# ZNF91 is an endogenous repressor of the molecular phenotype associated with X-linked dystonia-parkinsonism (XDP)

**DOI:** 10.1101/2023.10.20.563263

**Authors:** Jimi L. Rosenkrantz, Sanaz Raghib, J. Elias Brandorff, Ashni Kapadia, Christine A. Vaine, D. Cristopher Bragg, Grace Farmiloe, Frank M.J. Jacobs

**Author notes:** Correspondence to: Frank M. J. Jacobs, Science Park 904, 1098 XH Amsterdam, The Netherlands.

## Abstract

**Background:** X-linked dystonia-parkinsonism (XDP) is a severe neurodegenerative disorder resulting from the insertion of an intronic SINE-Alu-VNTR (SVA) retrotransposon in the *TAF1* gene. Recent research has revealed that the pathogenic XDP-SVA insertion leads to dysregulation of *TAF1* transcription, including increased intron retention and decreased expression of exons surrounding the insertion. The Krüppel-associated box (KRAB) zinc finger protein, ZNF91, is a critical repressor of SVA retrotransposons. However, it remains unclear whether ZNF91 is able to repress the XDP-SVA insertion and how this influences the XDP-associated molecular phenotype. In this study, we investigate the role of ZNF91 in repressing the XDP-SVA insertion and its impact on the molecular phenotype associated with XDP.

**Methods:** Here, we used CRISPR/Cas9 to genetically delete ZNF91 in induced pluripotent stem cell (iPSC) lines derived from XDP patients, as well as isogenic control iPSC lines that lack the XDP-SVA insertion. Total RNA sequencing and capture RNA-sequencing were used to confirm ZNF91 deletion and to assess *TAF1* transcriptional changes between conditions. Furthermore, publicly available transcriptomic data from whole blood and different brain regions were used to assess ZNF91 expression levels across ages.

**Results:** We found that genetic deletion of ZNF91 exacerbates the molecular phenotype associated with the XDP-SVA insertion in patient cells, while no difference was observed when ZNF91 was deleted from isogenic control cells. Additionally, we observed a significant age-related reduction in ZNF91 expression in whole blood and brain, indicating a potential role of ZNF91 in the age-dependent onset of XDP.

**Conclusions:** These findings indicate that ZNF91 plays a crucial role in controlling the molecular phenotype associated with XDP. Since ZNF91 is a critical epigenetic repressor of SVAs, this suggests that epigenetic silencing of the XDP-SVA minimizes the severity of the molecular phenotype. Our results showing that ZNF91 expression levels significantly decrease with age provide a potential explanation for the age-related progressive neurodegenerative character of XDP. Collectively, our study provides important insights into the protective role of ZNF91 in XDP pathogenesis and suggests that modulating ZNF91 levels or targeted repression of the XDP-SVA could be novel therapeutic strategies worth exploring.

## Background

X-linked dystonia-parkinsonism (XDP; MIM:314250), also known as DYT3 or Lubag syndrome, is a rare, progressive neurodegenerative disorder that primarily affects adult males with maternal ancestry from the Philippine Island of Panay. In the local Filipino dialect, XDP is referred to as “lubag,” meaning “twisted,” due to its characteristic dystonia (abnormal muscle tone and movements) and Parkinsonism (tremors, rigidity, and difficulty with movement) [1,2]. XDP was first described in the clinic nearly 50 years ago [3]. Since then, extensive research has revealed that it is caused by a 2.6 kb SINE-Alu-VNTR (SVA) retrotransposon insertion in intron 32 of the TATA-binding protein-associated factor-1 gene (*TAF1*) on chromosome Xq13 [4,5]. *TAF1* encodes a central protein of the transcription factor II D (TFIID) complex, which plays a crucial role in the transcription of genes involved in neurodevelopmental processes [6–11]. Recent studies using XDP patient cells with and without the disease-specific SVA insertion (XDP-SVA) have shown that XDP-SVA leads to dysregulation of *TAF1* transcription, including increased intron 32 retention and decreased expression of exons surrounding the XDP-SVA insertion [5,12]. *TAF1* dysregulation is thought to underlie the development of XDP, however the exact mechanism by which the XDP-SVA insertion contributes to *TAF1* dysregulation remains poorly understood.

SVA elements are a class of retrotransposons found exclusively in primate genomes [13] and belong to one of three active transposable element (TE) families in the human genome [14–16]. SVAs are a composite of several different repeat elements and consist of a (CCCTCT)n hexamer repeat, an Alu-like element, a variable number tandem repeat (VNTR), SINE-R, and Poly-A region. There are six subfamilies of SVAs (SVA_A to SVA_F). The two youngest subfamilies of SVAs, SVA_E and SVA_F, are human-specific and include many insertions that are polymorphic among human populations [13,17,18]. Polymorphic SVAs contribute to human genetic diversity and, in some cases, can lead to variation in traits and disease susceptibility [19–21]. This is in part due to the nature of SVA sequences, which possess robust gene regulatory potential and can exert a significant influence on nearby gene expression [22–24]. The epigenetic repression of SVAs is crucial in preventing their reinsertion into the genome but also to minimize the potential gene regulatory effects that may arise from their insertion. Many families of TEs, including SVAs, are silenced by Krüppel-associated box (KRAB) zinc finger (KZNF) proteins, which have been shown to recruit repressive epigenetic modifiers [22,25,26]. Recent findings have highlighted the critical role of the primate-specific zinc finger 91 (ZNF91) protein in repressing SVA elements [22,24]. These previous findings strongly suggest that ZNF91 binds and represses the pathogenic XDP-SVA insertion. However, the influence of ZNF91-mediated epigenetic repression on the molecular phenotype of XDP remains unclear.

In this study, we aimed to investigate how ZNF91 influences the XDP-SVA insertion and the associated molecular phenotype. Using CRISPR/Cas9 we genetically deleted ZNF91 from XDP induced pluripotent stem cell (iPSC) lines to assess the impact of ZNF91 on the XDP molecular phenotype. Our findings contribute important insights into the functional role of ZNF91 in the context of XDP and offer promising prospects for novel therapeutic approaches in combating XDP.

## Methods

### XDP iPSC culture

Parental XDP (MIN15i-33363.D and MIN04i-33109.2B) and dSVA (MIN31i-33363.D.3C2 and MIN32i-33109.2B.3A12) iPSC lines were obtained from the WiCell (**Supplemental Table 1**). XDP probands and respective dSVA iPSCs were cultured at 37 °C with 5% CO2 on Growth Factor Reduced Matrigel-coated (Corning) plates in mTeSR1 (STEMCELL). Media was changed daily, and the cells were split when they reached 80-90% confluency with Accutase. After each passage, 10 μM Y-27632 Rock inhibitor (Tocris) was added to the media and removed after 24 hours by refreshing the media.

### CRISPR‒Cas9 genetic deletion of ZNF91

For CRISPR/Cas9 deletion of ZNF91, two gRNAs designed to target exon 1 of ZNF91 [24] were cloned and inserted into pX330-SpCas9-HF1 (Addgene #108301). In addition to the gRNAs, pCAG. GFP (Addgene #11150) was cotransfected for FACS sorting of iPSCs. A day before transfection, iPSCs were seeded in 60 mm dishes coated with GFR Matrigel and incubated in mTESR1 with 10 μM Y-27632 Rock inhibitor (Tocris). At the time of transfection, the cells were 40-60% confluent, and their media was replaced one hour before transfection with prewarmed mTeSR1 and ROCK inhibitor. For each of the 60 mm dishes, 1.4 μg of each ZNF91-gRNA and 0.2 mg of pCAG. GFP plasmids were transfected. First, 125 µl optiMEM (Thermo Fisher) and 12.5 µl Lipofectamine stem (Thermo Fisher) were mixed, the plasmids were combined, and optiMEM was added to adjust the volume with the Lipofectamine mixture. The plasmids and Lipofectamine were mixed and incubated for 10 minutes at room temperature before adding the lipid-DNA mix to the cultures. After transfection, the cells were incubated for 24 hours. The media was changed the next day with prewarmed mTeSR1, and the cells were maintained for an additional 24 hours in the same cell culture condition. For sorting GFP-positive cells, the cells were supplemented with 10 μM Rock inhibitor an hour before Fluorescence-activated cell sorting (FACS). For collecting single cells, the plates were washed twice with pre-warmed PBS without Ca2+ and Mg2+ (Phosphate-buffered saline), subsequently Versene Solution was added to the plates and incubated at 37 °C for 7 minutes. The cells were filtered through a 35-mm mesh cell strainer and sorted on a FACS Aria III with a 100 μm nozzle. Sorted cells were seeded at clonal density of 6-8×10^3^ cells per 9.5cm^2^ dishes coated with GFR Matrigel and supplemented by the Rock inhibitor (10 μM) and mTeSR1. In addition, extra single cells (1-3×10^3^ cells per 9.5cm^2^ dishes) were collected in MEF (Mouse Embryonic Fibroblasts) supplemented with the Rock inhibitor (10 μM), Fibroblast Growth Factors (FGF, 8 ng/ml) and ESC medium. The media was changed the next day and adding the 10 μM Y-27632 Rock inhibitor (Tocris) was continued for two more days. Single colonies were manually scooped into separated wells of 96-well plates. Propagation of the cells was done by diluting Versene Solution with PBS (1:3) to use in genotyping, culturing, and freezing.

Genotyping of the expanded cells was done when the cells reached 90-95% confluency. The cells were treated with a lysis buffer (10 mM Tris-Cl pH 8.0, 10 mM disodium-EDTA, 10 mM NaCl, 0.5% w/v sarcosyl, 40 μg/ml proteinase K) overnight and incubated at 60 °C. The following day, after precipitating DNA with cold 95% ethanol supplemented with 75 mM NaCl and washing with 70% ethanol, genomic DNA (gDNA) was extracted and used for PCR analysis. Four primer sets, targeted internal and external deletion sites of ZNF91, were designed and used for the detection of successful excision. After selecting ZNF91 knock-out and Wild-type colonies based on the PCR amplicon sizes, the selected colonies were sent to Sanger sequencing for confirmation of the interested gene deletion.

### Mapping of ZNF91 ChIP-seq data to the XDP-SVA insertion

Publicly available ZNF91 ChIP-seq data from hESCs [24] was mapped to the XDP BAC clone (AB191243.1). First, raw fastq sequencing reads were trimmed of poor quality and adapter sequences using trimmomatic [27] (trimmomatic PE -phred33 ILLUMINACLIP:TruSeq3-PE.fa:2:30:10 LEADING:3 TRAILING:3 SLIDINGWINDOW:4:15 MINLEN:40). Next, the trimmed reads were aligned to the XDP BAC clone using BWA-MEM aligner using default settings [28]. Resulting SAM files was converted into sorted BAM file using samtools [29]. For visualization, BAM files were converted into bigwig files using deeptools bamCoverage [30] (bamCoverage --outFileFormat bigwig --binSize 1 --normalizeUsing None -- minMappingQuality 1). Bigwig ZNF91 ChIP-seq coverage results were visualized along the XDP BAC clone using the Integrative Genomics Viewer (IGV) desktop application [31].

### RNA-sequencing

#### Total RNA sequencing

Three Sanger-validated ZNF91 KO and WT clones per condition were sent for RNA sequencing. For this, cells were washed once with PBS before collection in TRIzol Reagent. Total RNA was extracted using TRIzol Reagent (Thermo Fisher), followed by gDNA removal and RNA purification using an RNA clean and concentrator kit (Zymo Research). Ribosomal RNA was depleted from total RNA with the rRNA depletion kit (NEB# E6310) and subsequently prepared for RNA-seq with the NEBNext Ultra Directional RNA Library Prep Kit (NEB #E7420) at GenomeScan. Samples were sequenced at 150 bp paired-end at an Illumina Hiseq 4000 device.

#### Capture RNA-sequencing (Cap-seq)

Two different sets of custom designed MyBaits probes (Arbor Bioscience) were used throughout this project. The first set of probes (MyBaits1) was designed to target and capture the *TAF1* cDNA sequence. The second set of probes (MyBaits2, Design ID: D10165*TAF1*) was designed to target and capture both cDNA and intron sequences of the *TAF1* gene. The first Cap-seq experiment (using the MyBaits1 probe set) was performed on a total of 12 iPSC samples from patient 1, including three replicates of a single clone from each cell line. The second Cap-seq experiment (using the MyBaits2 probe set) was performed on a total of 24 iPSC samples from patient 1 and patient 2, including three replicates of a single clone from each cell line.

TRIzol RNA isolation was performed as described above. Ribosomal depleted RNA-seq libraries were prepared using the TruSeq® Stranded Total RNA Library Prep Kit Human/Mouse/Rat kit (Illumina, 20020596) following manufacturer’s instructions. Libraries were quantified using DNA HS Bioanalyzer chip, and equal molar amounts were pooled together. Custom MyBaits probes kit was used according to the manufacturer’s instructions to capture and enrich pooled sequencing libraries. Briefly, libraries were blocked using Block O, C, and X buffers and then hybridized with capture probes for 16-24 hours at 65C. Samples were washed and cleaned, and KAPA HiFi HotStartReadMix was used to PCR amplify (14 cycles) the libraries. The final amplified and enriched libraries were cleaned using AmPureXP beads and sequenced on Illumina NextSeq500 using Mid Output flow cell 2×75 cycles.

### Analysis of RNA-sequencing data

Raw fastq sequencing reads were trimmed of poor quality and adapter sequences using FASTP [32] (fastp --qualified_quality_phred 30). Trimmed fastq files were then mapped to UCSC hg38 human reference genome using STAR [33] (STAR –outFilterMultimapNmax 20 -- outFilterMismatchNmax 5 --alignMatesGapMax 0 --alignIntronMax 0 -- outFilterMatchNminOverLread 0.66 --alignEndsType EndToEnd). The human gene annotation file (Homo_sapiens.GRCh38.104.gtf) was downloaded from ENSEMBL and converted to UCSC hg38 compatible annotations using the chromToUcsc python script. The final gene annotation file used also included an annotation for the intronic region upstream of the XDP-SVA insertion (*TAF1*_partial*TAF1*_intron32, chrX:71424239-71440488). The STAR-mapped BAM files and our custom gene annotation file were used to generate both transcript-level and exon-level count tables using FeatureCounts [34] (featureCounts -M -t exon). Count tables were filtered to only include transcript and exon IDs targeted by MyBaits probes. Normalization and differential expression analysis were performed using DEseq2 [35]. For both Cap-seq experiments expression was normalized to *TAF1* exons 1-28, and differential expression was performed using default DEseq2 parameters. A transcript of exon was considered differentially expressed if padj<0.05 and change was greater than 25% (L2FC>0.3219 or L2FC<-0.4150).

### Intron 32 splicing analysis

Data from both Cap-seq experiments (MyBaits1 and MyBaits2) were used for intron retention analysis. The number of reads mapping to the last bp of *TAF1* exon 32 (chrX:71424238), and the first bp of intron 32 (chrX:71424239) were extracted from indexed bam files using the igvtool command line tool [31] (igvtools count -w 1 –query). The number of reads spliced from exon 32 to exon 33 were extracted from bam files using samtools [29] by searching the CIGAR string for 29931bp splicing (’29931N’) using the following command (samtools view -F 0×4 $BAM chrX:71424229-71424248 | cut -f 6 | grep ’29931N’ | wc -l). The proportion of *TAF1* transcripts with canonical splicing from exon 32 to exon 33 was calculated by dividing the number of reads spliced from exon 32 to exon 33 by the number of reads mapping to the last bp of *TAF1* exon 32 (chrX:71424238). The proportion of *TAF1* transcripts with intron 32 retention was calculated by dividing the number of reads mapping to the first bp of intron 32 (chrX:71424239) by the number of reads mapping to the last bp of *TAF1* exon 32 (chrX:71424238).

### Analysis of ZNF91 expression across age

The whole blood microarray expression data used for the analyses described in this manuscript were obtained from The 10,000 Immunomes Project (10Kimmunomes) [36] (https://comphealth.ucsf.edu/app_direct_i/10kimmunomes/) on 06/09/23. The brain RNA-seq expression data, from thirteen distinct regions of the brain, were obtained from V8 of The Genotype-Tissue Expression (GTEx) project [37] (gtex_analysis_v8, rna_seq_data, GTEx_Analysis_2017-06-05_v8_RNASeQCv1.1.9_gene_tpm.gct) on 06/09/23. For the GTEx data, preprocessing involved merging tissue files with subject phenotypes allowing us to match gene expression levels with age for each subject. It’s important to note that the GTEx database provides subjects’ ages in a ten-year range; for this analysis, we chose to use the lower limit of each range as the representative age. For instance, a subject classified within the 50-59 years old range by GTEx would be considered 50 years old for the analysis. For the gene expression analysis, we investigated the ZNF91 gene along with *TAF1* and a random selection of 18 other ZNF genes from the GTEx database.

Our analysis of gene expression relied on the following Python libraries: pandas, statsmodels, and matplotlib for data processing, statistical modeling, and visualization, respectively. First, the dataset was imported using pandas and two subsets were defined: ’Age’ (the independent variable, x) and ’Expression’ (the dependent variable, y, indicating gene expression). Next, the statsmodels function ’add_constant’ was applied to the ’Age’ variable, introducing a constant to the independent variable. This step is important for the ordinary least squares (OLS) regression model (https://www.statsmodels.org/stable/index.html) since it estimates the intercept of the regression line. Following the creation of the model via the sm.OLS function, we fit the model to our data and generated a comprehensive statistical summary. This summary included two critical metrics: the adjusted R-squared value and the p-value. A p-value less than 0.05 was considered statistically significant. Matplotlib was employed to generate scatterplots with fitted regression lines and heatmaps (https://matplotlib.org/stable/users/index.html).

## Results

### Generation of stable ZNF91 knockout XDP iPSC lines

The pathogenic XDP-SVA insertion is situated within intron 32 of the *TAF1* gene. It is a full-length SVA element comprising the characteristic (CCCTCT)n hexamer repeat, an Alu-like region, a VNTR, SINE-R, and a Poly-A region [13,38] (**Fig. 1A**). SVAs, like many TE families, are silenced through the recruitment of members of the KRAB zinc finger (KZNF) family. The KZNF recruits KAP1 [39,40], along with repressive epigenetic modifiers, which repress the TE [41–43]. Our research group has recently demonstrated that the ZNF91 KZNF protein serves as a major repressor of SVA retrotransposons within the human genome, accomplishing repression by binding to both the Alu-VNTR and VNTR-SINE border regions within SVA elements [24]. Given that the XDP-SVA insertion contains intact Alu, VNTR, and SINE regions, it is expected that ZNF91 will bind and repress this specific SVA element. To visualize the predicted binding sites of ZNF91 at the XDP-SVA element, we mapped ZNF91 ChIP-seq data from human embryonic stem cells (hESCs) [24] to the XDP-SVA insertion. The analysis revealed a prominent peak at the Alu-VNTR border region, indicating that this region is likely bound and repressed by ZNF91 within the pathogenic XDP-SVA insertion (**Fig. 1A**). This finding is in accordance with prior research by Haring et al., which similarly demonstrated ZNF91 binding at this specific region within SVA elements in the human reference genome [24].

**Figure 1.**
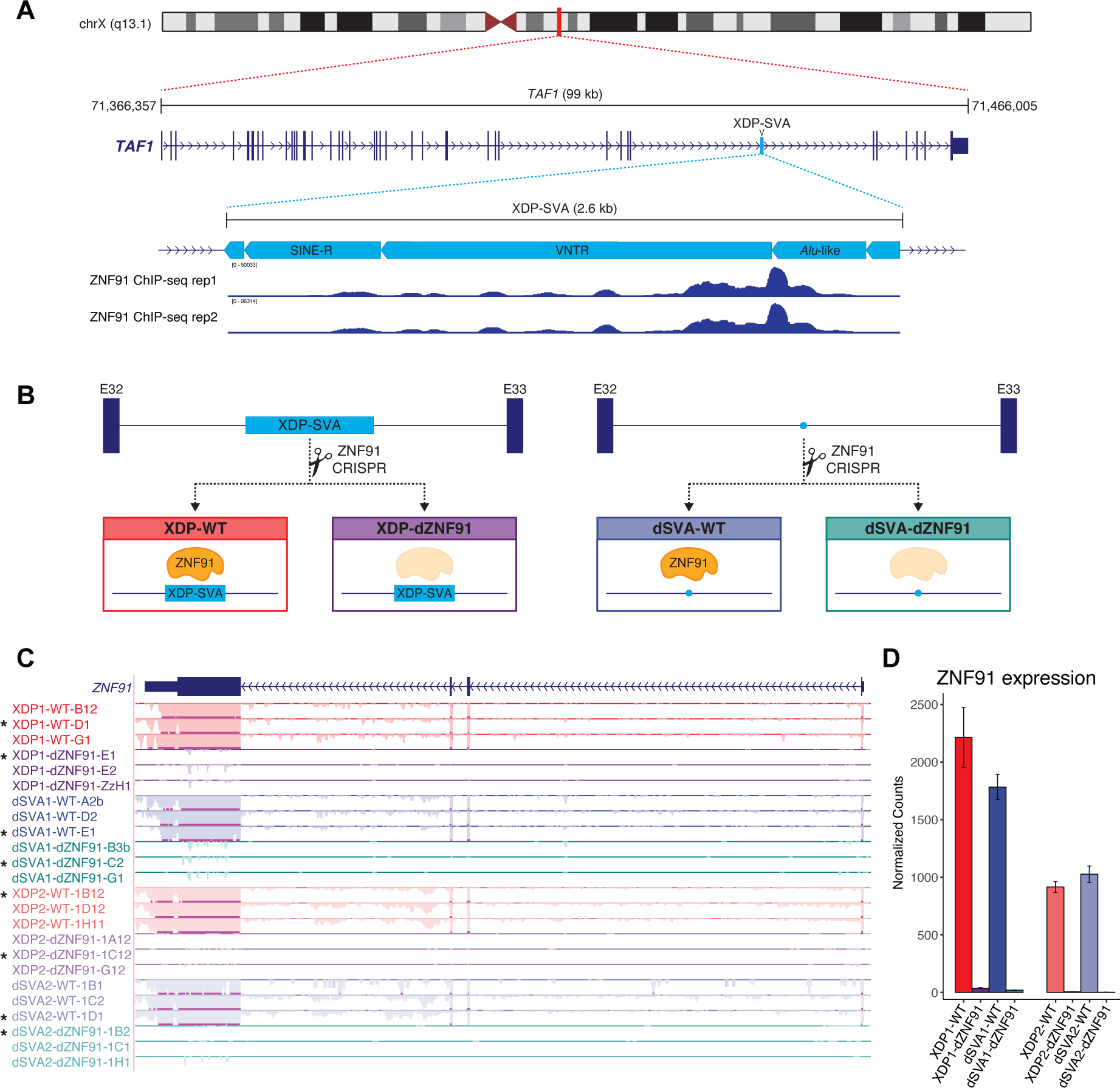
Generation of stable ZNF91 knockout XDP iPSC lines. (A) Schematic of TAF1, XDP-SVA insertion, and ZNF91 ChIP-seq signal. (Top) Ideogram of human Chromosome X, highlighting the genomic location of the TAF1 gene. (Middle) Detailed view of the TAF1 gene structure, the XDP-SVA insertion within intron 32 is highlighted in cyan. (Bottom) Schematic depiction of the XDP-SVA insertion, showing hESC ZNF91 ChIP-seq signal over the distinct subregions of the XDP-SVA element. (B) Schematic of CRISPR/Cas9 ZNF91 knockout in XDP and dSVA iPSCs. (C) Total RNA-seq coverage tracks at the ZNF91 locus highlighting loss of ZNF91 gene expression in ZNF91 knockout cell lines. Tracks are scaled to total library size. *Denotes samples used for capture RNA-seq analysis. (D) Quantification of ZNF91 gene expression. Each bar represents the mean ± SEM of N=3 replicate clones of each condition. DEseq2 default parameters were used for normalization.

It was recently established that the presence of the XDP-SVA insertion leads to dysregulation of *TAF1* transcription [5,12]. However, it remains unclear how binding of a repressive KZNF protein to this element influences the XDP molecular phenotype. To assess the functional implications of ZNF91 binding and repression on the XDP-SVA insertion and associated molecular phenotype we used CRISPR/Cas9 to genetically delete *ZNF91* in iPSCs derived from two XDP patients (XDP1 and XDP2) [44] and corresponding isogenic control iPSCs with XDP-SVA excised (dSVA1 and dSVA2) [5] (**Supplemental Table 1**). For each cell line, three *ZNF91* knockout (KO) and wild-type (WT) clones were established (**Fig. 1B, Supplemental Table 1**). To confirm the successful editing of *ZNF91* expression, total RNA-seq analysis was performed, revealing the absence of *ZNF91* expression in all *ZNF91* KO (dZNF91) clones and its presence in all *ZNF91* WT clones (**Fig. 1C-D, Supplemental Table 2**). The successful genetic deletion of the *ZNF91* gene in XDP iPSCs provides a valuable model to investigate the functional consequences of ZNF91 binding and repression on the XDP-SVA insertion and its associated molecular phenotype.

### Knockout of ZNF91 exacerbates the XDP molecular phenotype

#### Deletion of ZNF91 increases retention of the *TAF1* intron that harbors the XDP-SVA insertion

The presence of the XDP-SVA insertion leads to an elevated level of *TAF1* intron 32 expression [5]. This phenomenon was further validated in our study through analysis of total RNA-seq data from patient 1 and 2 iPSCs, which showed a marked increase in *TAF1* intron 32 expression in XDP compared to dSVA isogenic control samples (**Fig. 2A, Supplemental Table 2**). To investigate whether KO of ZNF91 affects this molecular phenotype, we developed a capture probe set with baits covering the canonical *TAF1* mRNA sequence and all introns of the *TAF1* gene and performed capture RNA sequencing (Cap-seq) on patient 1 and 2 cell lines. Differential expression (DE) analysis of ZNF91 KO compared with WT patient cells revealed a significant 130-250% upregulation of intron 32 expression in both patient cell lines (XDP1-dZNF91 vs. XDP1-WT, padj=1.37E-5, L2FC=1.22; XDP2-dZNF91 vs. XDP2-WT, padj=7.04E-13, L2FC=1.81) (**Fig. 2B-C, Supplemental Table 3**). Furthermore, analysis of intron 32 splicing showed that canonical splicing was decreased by ∼13% and ∼32%, while intron 32 retention was increased by ∼10% and ∼29% in ZNF91 KO cells compared with WT cells for patients 1 and 2, respectively (**Fig. 2D**). DE analysis revealed no significant change in intron 32 expression level when ZNF91 was deleted from the dSVA isogenic control cell lines (dSVA1-dZNF91 vs. dSVA1-WT, padj=0.9670, L2FC=0.06; dSVA2-dZNF91 vs. dSVA2-WT, padj=0.8754, L2FC=-0.07) (**Fig. 2B-C, Supplemental Table 3**) and negligible differences in canonical intron 32 splicing and retention levels were observed in ZNF91 KO compared with WT isogenic control cells (**Fig. 2D**). Similar results were obtained for patient 1 from a pilot Cap-seq experiment (excluding intron probes) (**Supplemental Fig. S1A-C, Supplemental Table 4**), providing further support that ZNF91 KO exacerbates the intron 32 molecular phenotype only in the presence of the XDP-SVA insertion and that increased expression of intron 32 is due in part to failed splicing and intron retention.

**Figure 2.**
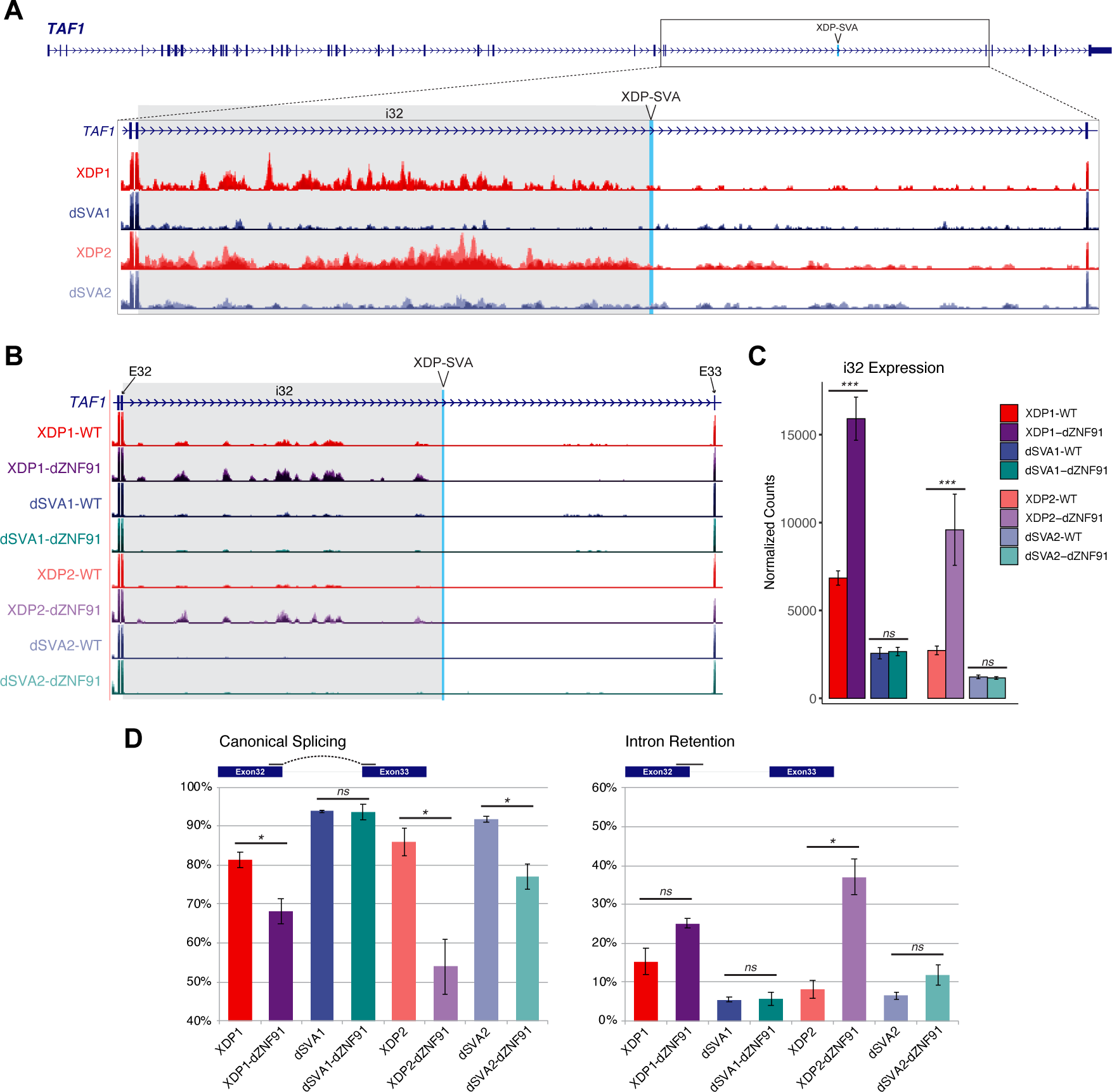
Deletion of ZNF91 increases retention of the TAF1 intron that harbors the XDP-SVA insertion. (A) Total RNA-seq coverage tracks at the TAF1 locus highlighting increase of intron 32 (i32) expression in XDP cells (red) compared to dSVA isogenic controls (blue). Tracks are scaled to TAF1 exons 1-28. (B) Capture RNA-seq coverage tracks at TAF1 intron 32, highlighting increased i32 expression in ZNF91 knockout XDP (purple) compared to WT XDP (red) cells. (C) Quantification of TAF1 i32 expression. A significant upregulation in ZNF91 KO XDP (purple) compared to WT XDP (red) cells is observed. Each bar represents the mean ± SEM of N=3 replicates (three replicates of single clones) and the DE analysis was performed using DESeq2 with default parameters (Wald test, corrected for multiple testing using the Benjamini and Hochberg method), except for normalization to TAF1 exons 1-28. Genes were considered differentially expressed if they had an adjusted p-value (padj) < 0.05 and exhibited greater than 25% change in expression (fold-change > 1.25 or <0.75). *padj<0.05, **padj<0.01, ***padj<0.001, ns = not significant. (D) TAF1 intron 32 splicing results. The percentage of reads with canonical splicing of intron 32 are displayed on the right, and the percentage of exon 32 reads with retained intron are displayed next on the left. Percentages are expressed as the mean ± SEM of three replicate samples. Statistical significance was assessed using a two-sided t-test, *p-value<0.05, ns = not significant.

#### Deletion of ZNF91 decreases transcription of *TAF1* exons located downstream of the XDP-SVA insertion

Previous studies have reported a slight decrease in the overall expression level of the *TAF1* gene due to the presence of the XDP-SVA insertion [5,12]. Subsequent analysis at the exon level further elucidated that the exons located in proximity to the XDP-SVA insertion site are primarily responsible for this observed decrease [5]. To investigate the impact of ZNF91 KO on this molecular phenotype, we utilized Cap-seq data and conducted exon-level differential expression analysis of the *TAF1* gene. In the case of patient 1, a significant reduction in the expression of downstream exons (exons 34-36 and 38) was observed upon deletion of ZNF91 in XDP1 iPSCs (XDP1-dZNF91 vs. XDP1-WT). However, no change in downstream exon expression was observed when ZNF91 was deleted in dSVA isogenic control cells (dSVA1-dZNF91 vs. dSVA1-WT) (**Fig. 3A-C, Supplemental Fig. S2, Supplemental Table 3-4**). Similar results were obtained for patient 2, where a significant reduction in the expression of all downstream exons (exons 33-38) was observed upon deletion of ZNF91 in XDP2 iPSCs (XDP2-dZNF91 vs. XDP2-WT). Again, no change in downstream exon expression was observed when ZNF91 was deleted in isogenic control dSVA2 iPSCs (dSVA2-dZNF91 vs. dSVA2-WT) (**Fig. 3A-C, Supplemental Table 3**). These findings provide compelling evidence that the loss of ZNF91 exacerbates the *TAF1* exonic molecular phenotype associated with XDP. Moreover, the absence of the effect in the dSVA isogenic control cells without XDP-SVA insertions underscores the specific regulatory impact of ZNF91 on the disease-causing XDP-SVA insertion.

**Figure 3.**
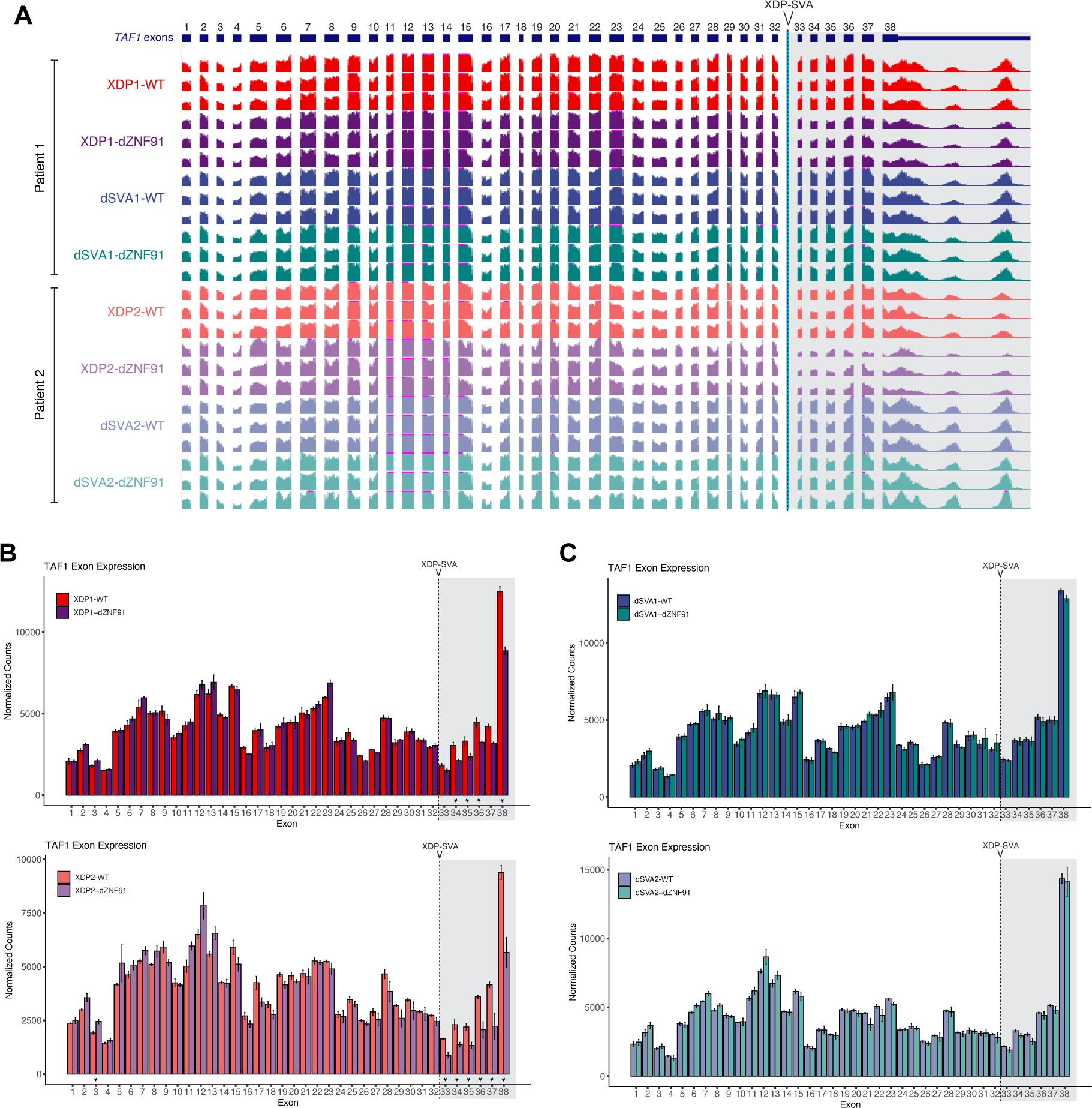
Deletion of ZNF91 decreases transcription of TAF1 exons located downstream of the XDP-SVA insertion. (A) Capture RNA-seq coverage tracks at TAF1 exons. A reduction of exon 33-38 expression is observed in ZNF91 knockout XDP (purple) compared to WT XDP (red) cells. Tracks are scaled to TAF1 exons 1-28. (B) Quantification of TAF1 exon 1-38 expression level in ZNF91 knockout XDP (purple) and WT XDP (red) cells. (C) Quantification of TAF1 exon 1-38 expression level in ZNF91 knockout dSVA (green) and WT dSVA (blue) cells. For panels B and C, each bar represents the mean ± SEM of three independent replicates (three replicates of single clones) and the DE analysis was performed using DESeq2 with default parameters (Wald test, corrected for multiple testing using the Benjamini and Hochberg method), except for normalization to TAF1 exons 1-28. Genes were considered differentially expressed if they had an adjusted p-value (padj) < 0.05 and exhibited greater than 25% change in expression (fold-change > 1.25 or < 0.75). *padj<0.05. For all panels, exons 33-38, downstream of the XDP-SVA insertion are highlighted with grey background.

### ZNF91 expression decreases with age

Despite the presence of the causative XDP-SVA mutation in all cells from birth, the onset of the XDP disease occurs later in life and specifically impacts certain regions of the brain [45–47]. The factors initiating the late onset and the brain-specific manifestations remain elusive. In our study, we provide compelling evidence that downregulation of ZNF91 expression exacerbates the molecular phenotype of XDP. Based on these findings, we hypothesize that an age-related reduction in *ZNF91* expression in the brain contributes to the disease onset. To investigate whether a general age-related reduction in ZNF91 expression exists in humans, we performed linear regression analysis on whole blood transcriptomic data from the 10K Immunomes Project [36]. To provide a single value summarizing both the magnitude and direction of the correlation, we devised a metric we call the Directional Coefficient of Determination (*dR^2^*). This metric is computed by multiplying the coefficient of determination (*R^2^*) by the sign of the correlation coefficient (*r*). It ranges from -1 to 1, with values closer to - 1 or 1 indicating stronger negative or positive relationships, respectively. Our analysis revealed a significant inverse correlation between *ZNF91* expression level and age (*dR^2^* = -0.062, p-value = 8.27E-06) (**Fig. 4A**). This result implies that approximately 6.2% of the variation in *ZNF91* expression can be explained by age in the regression model. This finding suggests that age plays a critical role in the regulation of *ZNF91* expression in blood, with older individuals exhibiting lower levels of *ZNF91* than younger individuals. To investigate the uniqueness of this observation for ZNF91, we extended our analysis to include 10 randomly selected ZNF genes. Intriguingly, none of these genes exhibited a significant inverse correlation with age (**Fig. 4B**), underscoring the distinct regulatory relationship between age and ZNF91 expression in the blood.

**Figure 4.**
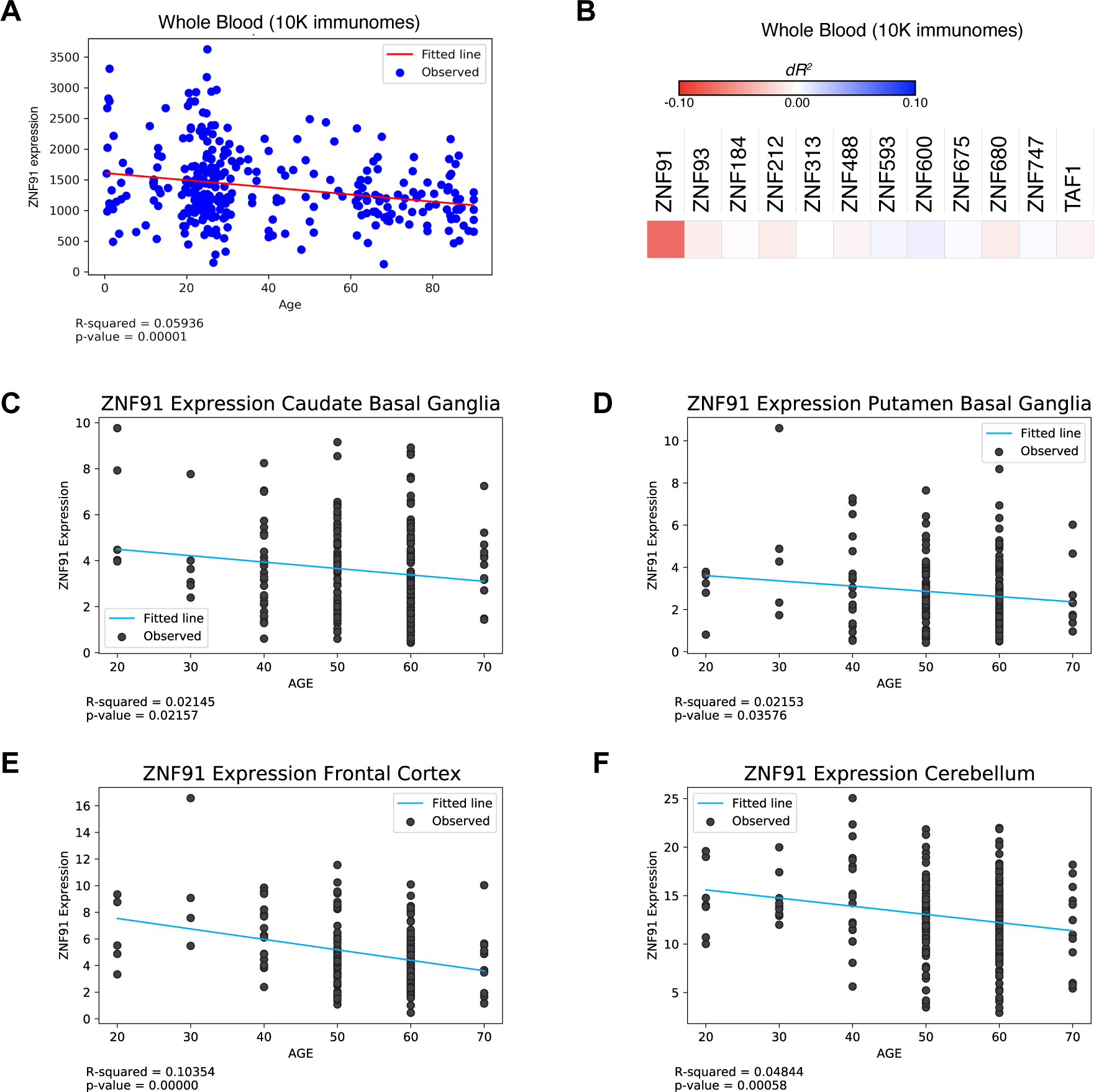
ZNF91 expression decreases with age. (A) Scatter plot illustrating the relationship between age (x-axis) and ZNF91 TPM expression level (y-axis) in whole blood samples from 311 individuals. Data sourced from the publicly available 10K Immunomes database. Linear regression R2 and significance level are included below plot. (B) Heatmap illustrating the linear regression results of gene expression level across age for 10 random ZNF genes in whole blood (10K immunomes data). Each box represents the directional R2 (dR2) value derived from the regression analysis, with the color intensity indicating the strength of the relationship. Red boxes signify an inverse correlation between ZNF91 expression and age, while blue boxes indicate a direct correlation. (C-F). Scatter plots illustrating the relationship between age and ZNF91 expression level in (C-D) striatal and (E-F) extra-striatal brain regions impacted in XDP. Data sourced from the publicly available GTEx database. Linear regression R2 and significance level are included below plot.

To investigate if the correlation between ZNF91 expression and age is also observed in brain regions implicated in XDP, we analyzed RNA-seq data from the GTEx database [37]. This encompassed 49 different tissues, of which 13 were distinct brain regions. A total of 32% of the tissues analyzed (16 out of 49) exhibited a significant inverse relationship between ZNF91 expression and age (**Supplemental Fig. S3**). This correlation was predominantly observed in brain tissues (12 out of 16), underscoring the specificity of the age-related reduction in *ZNF91* expression in the brain. Notably, a significant inverse correlation was observed in the striatal regions impacted by XDP, including the Caudate Basal Ganglia (*dR^2^* = -0.02145, p-value = 0.02157) and the Putamen Basal Ganglia (*dR^2^*= -0.02153, p-value = 0.03576) [48,49] (**Fig. 4C-D**). The correlation was even more pronounced in the frontal cortex (*dR^2^*= -0.10354, p-value = 2.02E-06) and cerebellum (*dR^2^*= -0.0484, p-value = 5.79E-04) - extrastriatal regions recently linked with structural thinning and volume loss in XDP patients [49,50] (**Fig. 4E-F**). These results indicate that age-related reductions in ZNF91 expression, especially within brain regions affected by XDP, may influence the disease’s onset or progression.

## Discussion

In this study, we investigated the impact of genetic deletion of ZNF91, a potent repressor of SVA elements, on the molecular phenotype associated with XDP. We demonstrated that knockout of ZNF91 aggravates the molecular phenotype, including an increase in *TAF1* intron 32 retention and a decrease in the expression of *TAF1* exons downstream of the XDP-SVA insertion. Additionally, our data show that there is a significant inverse correlation between age and ZNF91 expression levels in both whole blood and the brain. The results of this study provide important insights into the functional role of ZNF91 in the context of XDP. By observing the aggravated molecular phenotype in the absence of ZNF91, we can conclude that ZNF91 is an endogenous repressor of the molecular phenotype associated with XDP. Further, the observation that ZNF91 expression levels significantly decrease with age supports its potential involvement in disease onset and/or progression, which is known to occur later in life [45]. In summary, the findings from our study support the following proposed model: the age-related decline in ZNF91 expression unleashes the full pathogenic potential of the XDP-SVA insertion, thereby driving the onset of the disease (**Fig. 5**).

**Figure 5.**
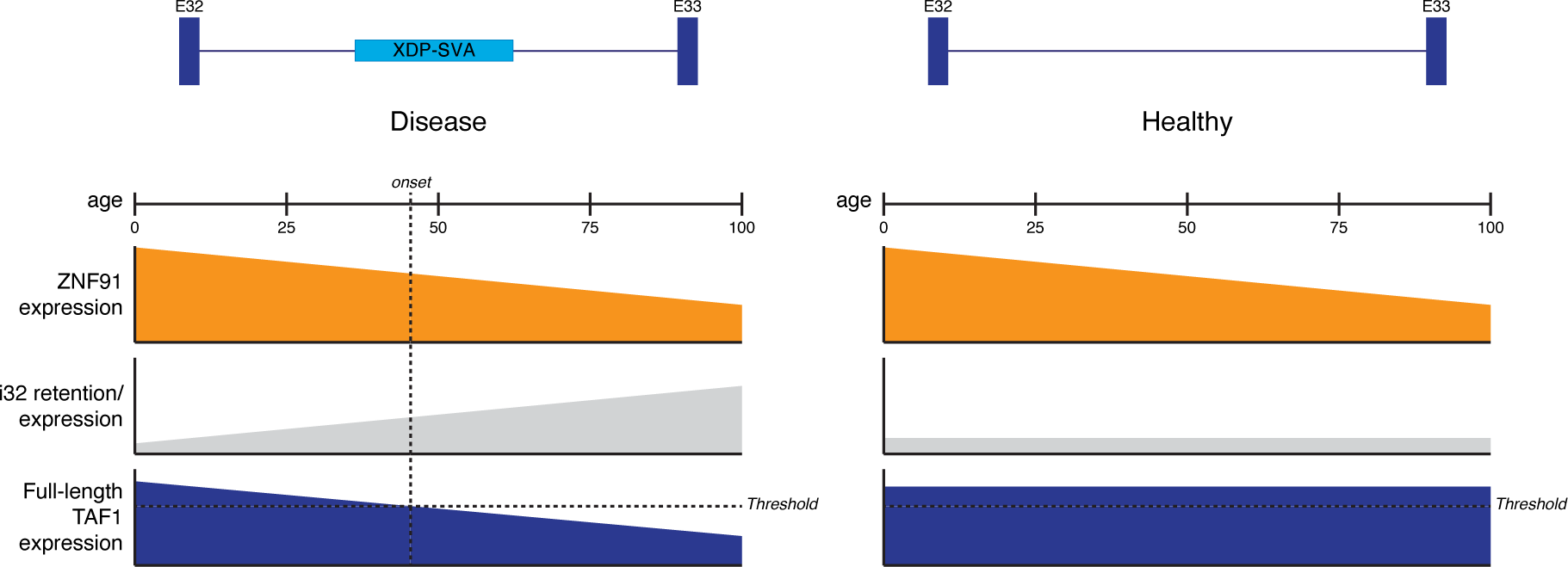
Proposed model: age-related reduction of ZNF91 unleashes full pathogenic potential of XDP-SVA. ZNF91 expression significantly decreases with age. Further, ZNF91 interacts specifically with the disease-specific XDP-SVA insertion and is an endogenous repressor of the associated molecular phenotypes. Thus, we propose that the age-related loss of ZNF91 leads to increased TAF1 intron 32 retention and decreased levels of full-length TAF1 transcripts in XDP patients. In contrast, the age-related loss of ZNF91 has no effect on TAF1 transcription in healthy individuals since they lack the XDP-SVA insertion.

This study represents a significant milestone as it is the first to successfully modulate *TAF1* dysregulation without altering any of the XDP-specific genetic variations [1,5,38]. Previous work utilizing CRISPR/Cas9 deletion of the XDP-SVA has been instrumental in dissecting the causal mechanism of XDP, revealing that the XDP-SVA insertion drives the XDP-specific *TAF1* transcriptional dysregulation [5,12]. The results of this study align with these previous findings and provide novel insights into how ZNF91 impacts the severity of the molecular phenotype. Given ZNF91’s established role as a repressor of SVAs, our results strongly suggest that epigenetic repression of the XDP-SVA by ZNF91 serves as a critical regulatory mechanism that keeps the molecular phenotype under control. This indicates a potential direct relationship between ZNF91-mediated epigenetic control and disease severity. These new findings provide valuable knowledge for understanding the underlying mechanisms of XDP and may contribute to the development of epigenetic-based therapies for this disorder.

Previous research has established the role of ZNF91 in binding and repressing the majority of SVAs in the human reference genome [22,24]. Our study builds upon this work and demonstrates the direct involvement of ZNF91 with a polymorphic disease-associated SVA. Moreover, our results highlight the influence of this interaction on the molecular phenotype of the disease. This underscores the importance of ZNF91 as a key regulator of SVA elements and its potential implications in the pathogenesis of XDP and other SVA-associated diseases. In addition to XDP, polymorphic SVA insertions have previously been associated with various other human diseases, including Lynch syndrome [51], hemophilia B [52], Fukuyama-type congenital muscular dystrophy (FCMD) [53], X-linked agammaglobulinemia (XLA) [54,55], autosomal recessive hypercholesterolemia (ARH) [56], and neurofibromatosis type I (NF1) [57]. Our findings may open new avenues of investigation into the mechanisms underlying these other disease-associated SVAs and their impact on disease susceptibility and progression.

The age of onset of XDP is highly variable (12-64 years old) [58], studies looking at the hexamer repeat length and nearby single nucleotide polymorphisms (SNPs) have succeeded in partially explaining this age-at-onset variability (50% and 13%, respectively) [4,58,59]. However, a large proportion (37%) of the age-at-onset variability is still unexplained. Given the findings of this study, it is reasonable to speculate that ZNF91 expression levels could contribute to the age-at-onset variability in XDP patients. Those with lower ZNF91 expression levels might experience more severe *TAF1* dysregulation, leading to an earlier onset of the disease. These epigenetic and transcriptomic differences go undetected by traditional DNA-based genome wide association studies (GWAS).

It is important to acknowledge that all experiments in this study were conducted using iPSCs, which primarily mimic early embryonic development—a stage seemingly unaffected by XDP. Consequently, the findings obtained from these cell types may not fully represent the events occurring in striatal medium spiny neurons (MSNs), which are predominantly affected by the disease [46]. Recent studies have reported the successful generation of MSNs from iPSCs [60–62]. The employment of iPSC-derived MSNs would provide more comprehensive insights into how ZNF91 deletion impacts the molecular phenotype in the specific neuronal cell types affected by XDP. However, fully recapitulating the affected cell types using any *in vitro* model poses challenges, considering that cell lines are limited to several months of culture, whereas the average age of onset for XDP is approximately 40 years old [45]. This discrepancy in “age” between *in vitro* models and the actual age of onset poses a difficulty in fully capturing the late-onset component of XDP *in vitro*. Despite these challenges, the use of iPSCs in this study provides valuable initial data and lays the groundwork for further research in this area. The combination of *in vitro* models with other approaches, such as animal models or postmortem human brain tissue analysis, may offer complementary insights into the complex disease mechanisms underlying XDP.

Previous investigators have proposed that guanine-rich sequences within the XDP-SVA might be susceptible to forming non-canonical DNA structures, called G-quadruplexes (G4) [4,63]. An intriguing hypothesis arising from our ZNF91 KO findings is that in the absence of ZNF91 binding and/or repression, the intronic XDP-SVA might be susceptible to the formation of G4 structures. As previous studies suggest, the presence of G4 structures within a gene can hinder RNA polymerase during transcription [64,65]. Blockage of RNA polymerase, and subsequent disruption of splicing, could explain the observed increase in intron 32 retention and subsequent decrease in downstream exon expression. While this hypothesis offers a compelling explanation for our results, additional investigations are essential to test this.

Several unanswered questions and intriguing directions for future research have emerged from this study, which hold the potential to further enhance our understanding of how ZNF91 expression may contribute to XDP pathogenesis. First, it remains unclear whether reduction of ZNF91 expression levels can trigger similar effects as its complete knockout and whether there is a “threshold” level of ZNF91 expression required for maximal repression of the phenotype. Investigating the consequences of varying ZNF91 expression levels, ideally in MSNs, would shed light on its dosage-dependent effects on the XDP-SVA insertion and the associated molecular phenotype. Additionally, it remains unclear whether the aggravated molecular phenotype observed upon loss of ZNF91 contributes to the onset or progression of this disease and whether the aggravation is caused simply by loss of ZNF91 binding or epigenetic derepression of the XDP-SVA insertion. Addressing these questions is important for determining whether modulation of ZNF91 expression and/or repression of the XDP-SVA are viable therapeutic approaches.

In conclusion, this study provides important insights into the functional role of ZNF91 in the context of XDP. The results demonstrate that ZNF91 deletion aggravates TAF1 dysregulation, indicating that ZNF91 acts as an endogenous repressor of the XDP-SVA and the associated molecular phenotype. Importantly, this study is the first to successfully modulate the XDP molecular phenotype without altering any XDP-specific genetic variants in the *TAF1* region. Our findings highlight the significance of ZNF91 as a key regulator of SVA elements and its potential implications in disease pathogenesis, not only in XDP but also in other SVA-associated disorders. In the broader context, our research contributes to the growing knowledge of intronic gene regulation in disease pathogenesis. Ultimately, the identification of ZNF91 as a potential therapeutic target in XDP and other SVA-associated diseases offers promising prospects for innovative approaches in disease treatment.

## List of abbreviations

ARH: Autosomal recessive hypercholesterolemia
Cap-seq: Capture RNA-sequencing
Cas9: CRISPR-associated protein 9
cDNA: Complementary DNA
ChIP-seq: Chromatin Immunoprecipitation Sequencing
CRISPR: Clustered regularly interspaced short palindromic repeats
DE: Differential expression
dSVA: XDP-SVA excised
DYT3: X-linked dystonia-parkinsonism (Alternative name)
ESC: Embryonic Stem Cell
FACS: Fluorescence-activated cell sorting
FCMD: Fukuyama-type congenital muscular dystrophy
FGF: Fibroblast Growth Factor
G4: G-quadruplexes
gDNA: Genomic DNA
gRNA: Guide RNA
hESCs: Human Embryonic Stem Cells
iPSC: Induced pluripotent stem cell
KAP1: KRAB-associated protein 1
KO: Knockout
KRAB: Krüppel-associated box
KZNF: KRAB zinc finger
L2FC: Log2 fold change
NF1: Neurofibromatosis type I
OLS: Ordinary least squares
padj: Adjusted P-value
PBS: Phosphate-Buffered Saline
PEI: Polyethylenimine
Poly-A: Poly-adenosine
RNA-seq: RNA-sequencing
SINE: Short Interspersed Nuclear Element
SVA: SINE-VNTR-Alu
TAF1: TATA-binding protein-associated factor-1
TE: Transposable Element
TFIID: Transcription Factor II D
VCM: Virus Containing Media
VNTR: Variable Number Tandem Repeat
WT: Wild type
XDP: X-linked dystonia-parkinsonism
XDP-SVA: XDP-specific SVA
XLA: X-linked agammaglobulinemia
ZNF91: Zinc Finger Protein 91

## Declarations

### Ethics approval and consent to participate

The present study involves the use of deidentified XDP patient iPSCs that were derived from a previous study [44]. In the primary study generating the iPSC lines used in this research [44], patients provided informed written consent for their biological samples to be used for scientific investigation.

### Consent for publication

All authors have given their consent for publication.

### Availability of data and materials

The datasets generated and/or analyzed during the current study are available in the NCBI Sequencing read archive (SRA; https://www.ncbi.nlm.nih.gov/sra/) repository under accession number PRJNA1004173. The ZNF91 ChIP-seq data was downloaded from NCBI Gene Expression Omnibus (GEO; https://www.ncbi.nlm.nih.gov/geo/) accession number GSE162571. The XDP BAC clone “Homo sapiens DNA, transposon SVA sequence” was downloaded from the NCBI Nucleotide database (https://www.ncbi.nlm.nih.gov/nuccore/AB191243.1/) accession number AB191243.1.

### Authors’ contributions

Conceptualization: FMJ and JLR; Methodology: JLR, SR, JEB, and AK; Validation: JLR, SR, and JEB; Formal Analysis: JLR and AK; Investigation: JLR, SR, and JEB; Resources: SR, CAV, and DCB; Writing – Original Draft: JLR; Writing – Review & Editing: JLR, DCB, GF, and FMJ; Visualization: JLR and AK; Supervision: FMJ and JLR; Project Administration: FMJ and JLR; Funding Acquisition: FMJ

## Acknowledgments

We would like to acknowledge the research subjects and clinical research staff of the Dystonia Partners Research Bank (Massachusetts General Hospital, Boston, USA) for their effort, foundational contributions to this work and commitment to improving the lives of people and families with XDP. Funding for this study was provided by the MGH Collaborative Center for X-Linked Dystonia-Parkinsonism (CCXDP). We would also like to thank Christopher Bragg, Amy Alessi, Johan Jakobsson, and all members of CCXDP for their valuable discussions and guidance on the project; all members of the Jacobs lab for their engaging discussions, and constructive feedback; Gonzalo Congrains Sotomayor for technical assistance with FACS; UvA sequencing core for technical assistance with Illumina sequencing; the Evolutionary Neurogenomics Group, the Molecular Neuroscience group, and others at the Swammerdam Institute for Life Sciences (SILS) for helpful discussions.

