## Supplemental Fig. for "ZNF91 is an endogenous repressor of the molecular phenotype associated with X-linked dystonia-parkinsonism (XDP)"

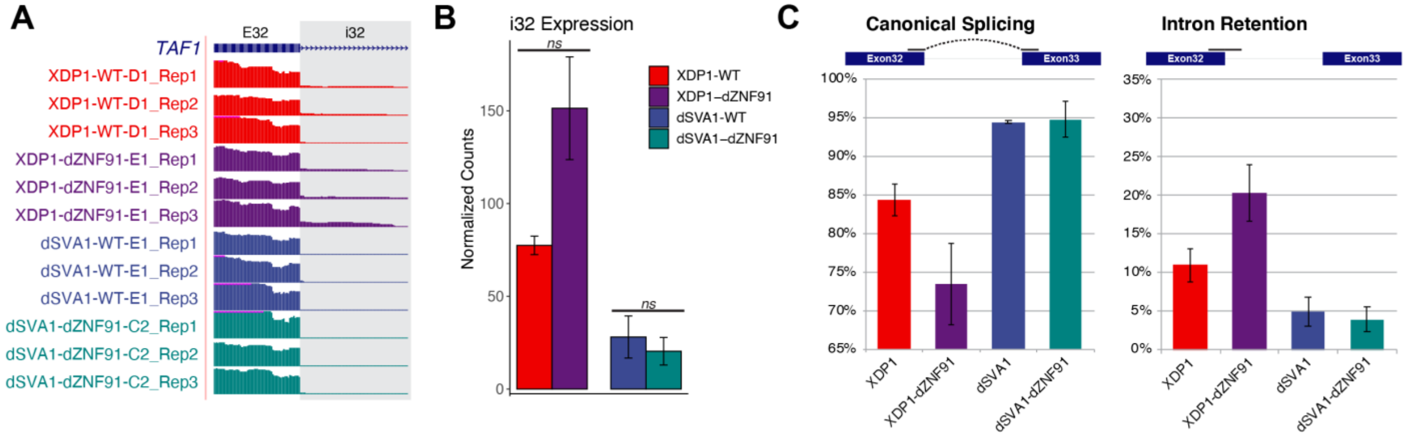


**Supplemental Figure S1. Deletion of ZNF91 increases retention of the TAF1 intron that harbors the XDP-SVA insertion (Pilot Capture RNA-seq results excluding intron probes).**
**(A)** Capture RNA-seq coverage tracks at TAF1 exon32/intron32 boundary, highlighting increased i32 expression in ZNF91 knockout XDP (purple) compared to WT XDP (red) cells. **(B)** Quantification of TAF1 i32 expression. A marked, but not statistically significant, increase of i32 in ZNF91 knockout XDP (purple) compared to WT XDP (red) cells is observed. Each bar represents the mean ± SEM of N=3 independent replicates (three replicates of single clones) and the DE analysis was performed using DESeq2 with default parameters (Wald test, corrected for multiple testing using the Benjamini and Hochberg method), except for normalization to TAF1 exons 1-28. Genes were considered differentially expressed if they had an adjusted p-value (padj) < 0.05 and exhibited greater than 25% change in expression (fold-change > 1.25 or <0.75). *padj<0.05, **padj<0.01, ***padj<0.001, ns = not significant. **(C)** TAF1 intron 32 splicing results. The percentage of reads with canonical splicing of intron 32 are displayed on the right, and the percentage of exon 32 reads with retained intron are displayed next on the left. Percentages are expressed as the mean ± SEM of three replicate samples. Statistical significance was assessed using a two-sided t-test, *p-value<0.05, ns = not significant.

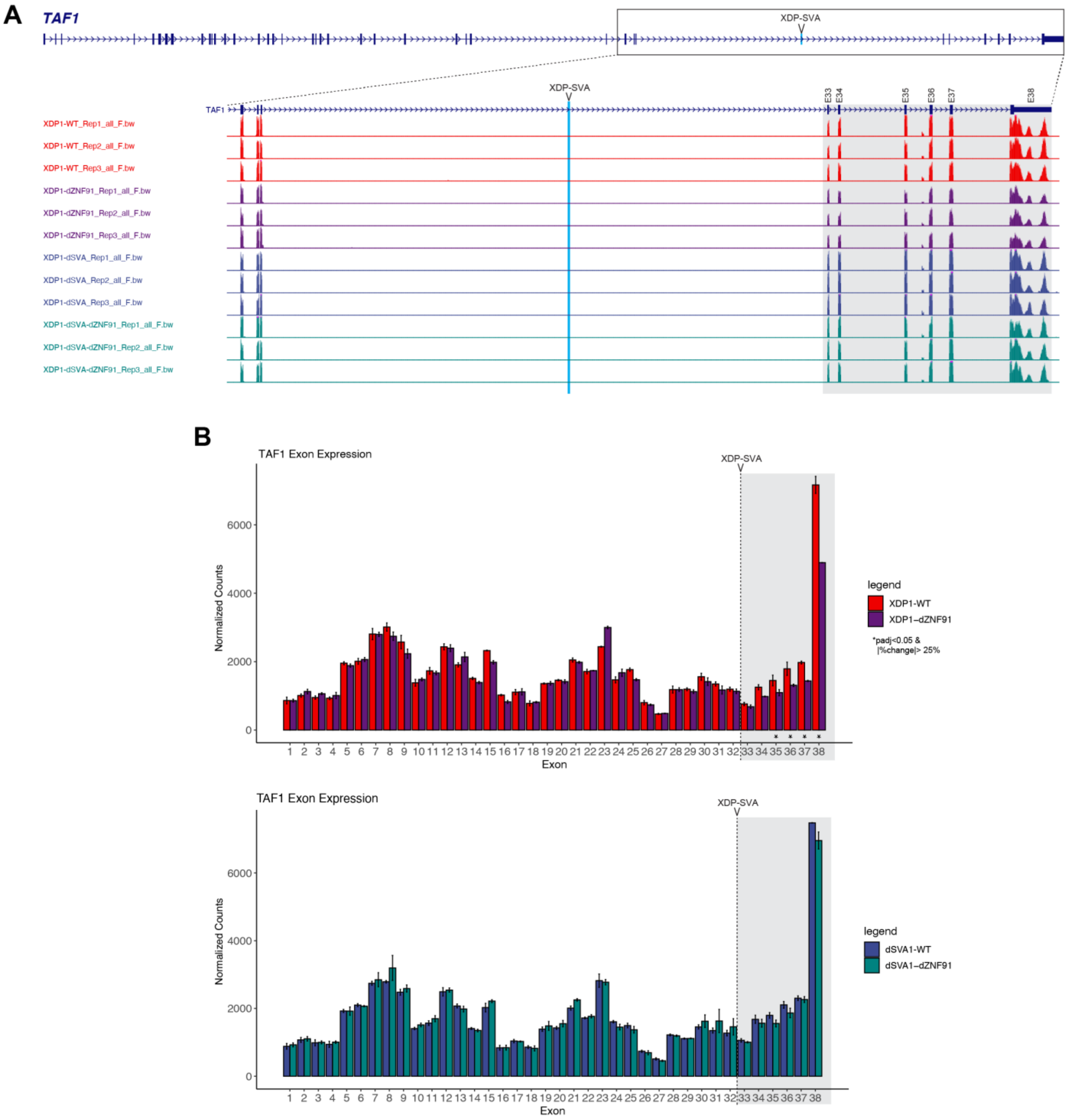


**Supplemental Figure S2. Deletion of ZNF91 decreases transcription of TAF1 exons located downstream of the XDP-SVA insertion (Pilot capture RNA-seq results excluding intron probes)**

**(A)** Capture RNA-seq coverage tracks at TAF1 exons. A reduction of exon 33-38 expression is observed in ZNF91 knockout XDP (purple) compared to WT XDP (red) cells. Tracks are scaled to TAF1 exons 1-28. **(B)** Quantification of TAF1 exon 1-38 expression level in ZNF91 knockout XDP (purple) and WT XDP (red) cells. **(C)** Quantification of TAF1 exon 1-38 expression level in ZNF91 knockout dSVA (green) and WT dSVA (blue) cells.

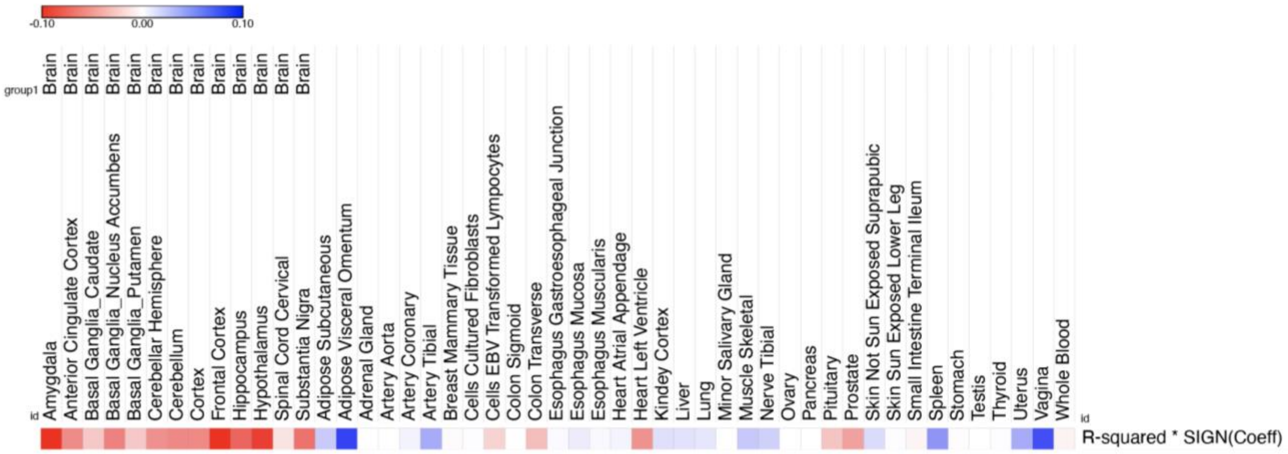


**Supplemental Figure S3. ZNF91 expression decreases with age specifically in the brain.**
Heatmap illustrating the linear regression results of ZNF91 expression across age for various GTEx tissues. Each box represents the directional $R^2$ ($dR^2$) value derived from the regression analysis, with the color intensity indicating the strength of the relationship. Red boxes signify an inverse correlation between ZNF91 expression and age, while blue boxes indicate a direct correlation. Only GTEx tissues with more than 30 samples were included in this analysis.
